# A recombinant fragment of Human surfactant protein D binds Spike protein and inhibits infectivity and replication of SARS-CoV-2 in clinical samples

**DOI:** 10.1101/2020.12.18.423415

**Authors:** Taruna Madan, Barnali Biswas, Praveen M. Varghese, Rambhadur Subedi, Hrishikesh Pandit, Susan Idicula-Thomas, Indra Kundu, Sheetalnath Rooge, Reshu Agarwal, Dinesh M. Tripathi, Savneet Kaur, Ekta Gupta, Sanjeev K. Gupta, Uday Kishore

## Abstract

**Rationale:** COVID-19 is an acute infectious disease caused by the Severe Acute Respiratory Syndrome Coronavirus 2 (SARS-CoV-2). Human surfactant protein D (SP-D) is known to interact with spike protein of SARS-CoV, but its immune-surveillance against SARS-CoV-2 is not known.

**Objective:** This study aimed to examine the potential of a recombinant fragment of human SP-D (rfhSP-D) as an inhibitor of replication and infection of SARS-CoV-2.

**Methods:** rfhSP-D interaction with spike protein of SARS-CoV-2 and hACE-2 receptor was predicted via docking analysis. The inhibition of interaction between spike protein and ACE-2 by rfhSP-D was confirmed using direct and indirect ELISA. The effect of rfhSP-D on replication and infectivity of SARS-CoV-2 from clinical samples was studied by measuring the expression of RdRp gene of the virus using qPCR.

**Measurements and Main Results:** *In-silico* interaction studies indicated that three amino acid residues in the RBD of spike of SARS-CoV-2 were commonly involved in interacting with rfhSP-D and ACE-2. Studies using clinical samples of SARS-CoV-2 positive cases (asymptomatic, n=7 and symptomatic, n=8 and negative controls n=15) demonstrated that treatment with 5μM rfhSP-D inhibited viral replication by ~5.5 fold and was more efficient than Remdesivir (100 μM). Approximately, a 2-fold reduction in viral infectivity was also observed after treatment with 5μM rfhSP-D.

**Conclusions:** These results conclusively demonstrate that the calcium independent rfhSP-D mediated inhibition of binding between the receptor binding domain of the S1 subunit of the SARS-CoV-2 spike protein and human ACE-2, its host cell receptor, and a significant reduction in SARS-CoV-2 infection and replication *in-vitro*.

## Introduction

The COVID-19 pandemic, caused by the Severe acute respiratory syndrome Coronavirus-2 (SARS-CoV-2) (1, 2), has affected ~ 58 million people across the globe and has claimed more than a million lives within its first year (3). The SARS-CoV-2 spike protein (S protein) is cleaved into S1 subunit, which is involved in host receptor binding, and S2 subunit, which is involved in membrane fusion, by the host’s transmembrane Serine Protease 2 (TMPRSS2) (4). This priming of the S protein by host proteases enables it to bind with the angiotensin-converting enzyme 2 (ACE2) receptor on the nasopharyngeal epithelial cells, leading to its entry into the host cell (4). While vaccines against the virus are being developed and trialled, the current therapeutic strategy is empirical and comprises of anti-viral medications and immunosuppressants (5).

The innate immune system plays a crucial role against SARS-CoV-2 infection; majority of infected individuals purge the virus within a few days with minimal involvement of adaptive immune response (6). Collectins are a group of humoral pattern recognition receptors, of which human lung surfactant protein D (SP-D), is known to act as a potent viral entry inhibitor, including HIV-1 and influenza A virus (7, 8). The primary structure of SP-D is characterised by an N-terminus that is involved in multimerization; a triple-helical collagenous region made up of Gly-X-Y repeats, an α-helical coiled-coil neck region, and a C-terminal C-type lectin or carbohydrate recognition domain (CRD) (9). The protective effects of SP-D against a range of bacterial, viral, and fungal pathogens leading to their agglutination, growth inhibition, enhanced phagocytosis, neutralisation, and modulation of immune responses are well documented (9, 10).

During the SARS-CoV epidemic in 2002, elevated levels of SP-D were reported in the serum of the patients infected with highly pathogenic β-CoV, SARS CoV (11). Purified SP-D has been shown to bind to the receptor-binding domain (RBD) of the glycosylated Spike protein of SARS-CoV, which shares 74% homology with the RBD of SARS-CoV-2 (12). In addition, SP-D also binds α-CoV, HCoV-229E, and inhibits infection in human bronchial epithelial cells (13). These mounting pieces of evidence encouraged exploration of the therapeutic potential of SP-D in COVID-19 patients.

In this study, we used a well-characterised recombinant fragment of human SP-D (rfhSP-D) comprising homotrimeric neck and CRD regions to study its protective effect against SARS-CoV-2 infection. As the recombinant form has the advantage of a smaller size to reach the distal lung locations and higher resistance to proteases and collagenases over the full-length SP-D, we evaluated the interaction of rfhSP-D with RBD and Spike of SARS-CoV-2 and its inhibitory potential against infection and replication of SARS-CoV-2 in clinical samples.

## Materials and Methods

### Clinical Samples

The clinical samples (Table 1), nasopharyngeal (NP) and oropharyngeal (OP) swabs (n=30) used in this study, were stored at the BSL-3 facility of the Institute of Liver and Biliary Sciences, Delhi. These clinical samples (n=15) were from symptomatic contact of lab-confirmed cases (Cat 2), hospitalised severe acute respiratory infections (SARI) case-patients (Cat 4), asymptomatic direct and high-risk contacts of lab-confirmed case (Cat 5a) and hospitalised symptomatic influenza-like illness (ILI) case-patients (Cat 6) that had tested positive by RT-PCR test for SARS-CoV-2. The samples obtained were placed in the viral transport medium (Hanks Balanced Salt Solution (HBSS) supplemented with 2% heat-inactivated FBS, 100 μg/ml Gentamicin and 0.5 μg/ml of Amphotericin B. NP and OP samples (n=15) that tested negative by RT-PCR test for SARS-CoV-2 were used as controls. The 50% Tissue culture Infective Dose (TCID_50_) of the clinical samples obtained was confirmed using an MTT assay. Briefly, 5 x 10^4^ Vero cells in Vero growth media (MEM Glutamax, supplemented with 10% Fetal Bovine Serum, 1% v/v Penicillin-Streptomycin and 1%v/v sodium pyruvate [Gibco, Thermofisher]) were seeded in a 96 well plate and grown overnight. The clinical samples from the 15 confirmed COVID-19 patients, and the 15 controls were added to the cells and incubated for 1h. Post incubation, the wells were washed with PBS twice, and fresh Vero growth medium was added to the cells. The cells were then incubated for 96 h at 37°C, 5% CO_2_. A 3-[4,5-dimethylthiazol-2-yl]-2,5-diphenyltetrazolium bromide (MTT) assay was performed to assess the viability of the cells by incubating the cells with 12mM of MTT for 4 h at 37°C, 5% CO_2_. The formazan created was dissolved using DMSO, and the samples were read at 590 nm using a microplate reader.

**Table 1:**
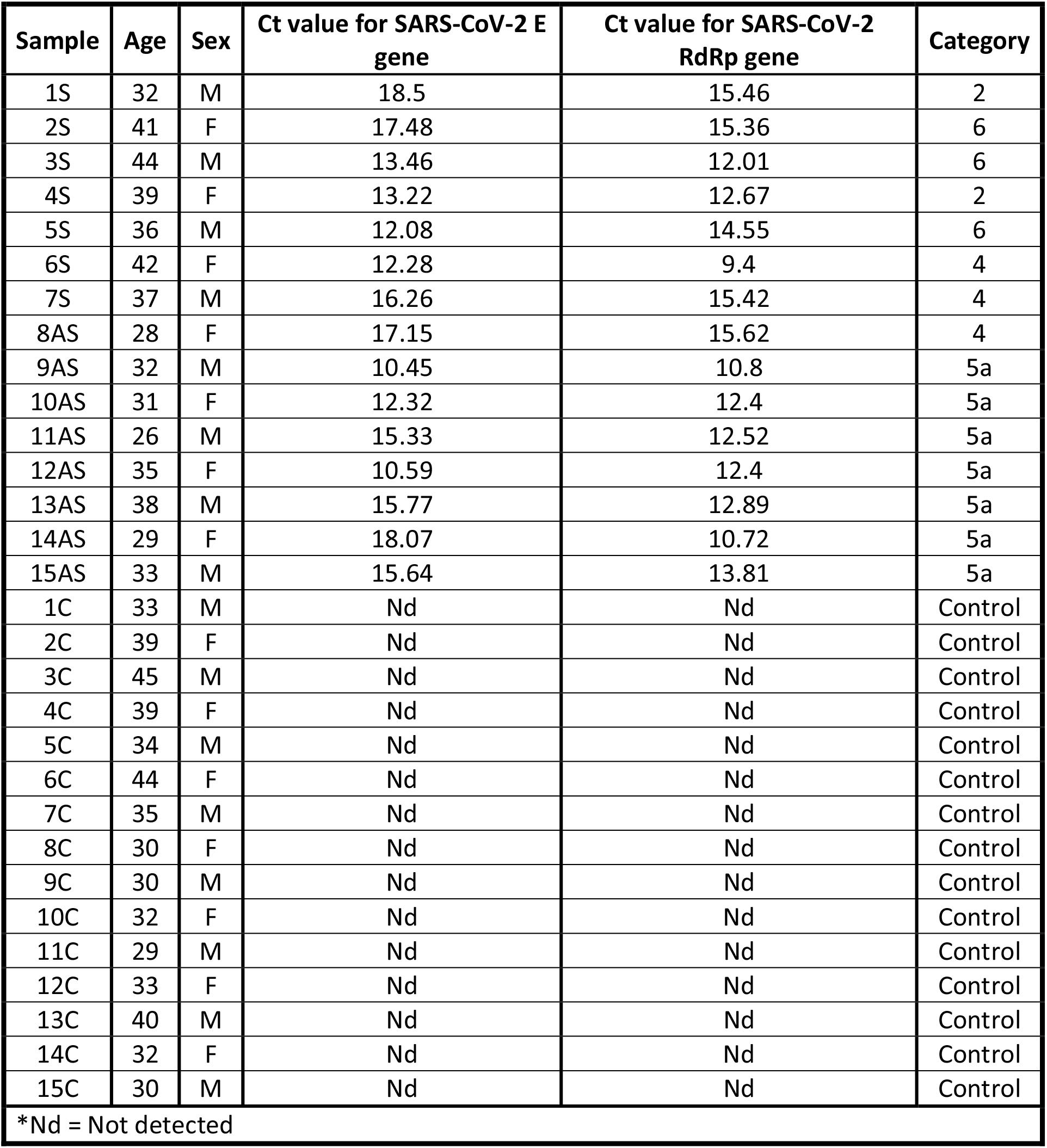
Characteristics of clinical samples utilised in the study

### *In silico* Analysis

The co-crystallised structure of human ACE-2 receptor with Spike S protein (PDB id: 6VW1) was separated into its receptor (ACE-2) and ligand (Spike S) components. The receptor and ligand were then re-docked using Patchdock web server (14, 15) to validate the docking protocol. The therapeutic agent, rfhSP-D trimer, (PDB id: 1PW9) was individually blind docked with the structure of RBD of S protein in the open conformation (PDB id: 6VYB) and dimeric ACE-2 (PDB id: 6VW1) using Patchdock. Top 100 docked poses were selected and further refined using FireDock web server (16, 17) for calculation of global free energy. The top 5 refined structures were filtered based on interactions between receptor binding motif (RBM) of S protein, CRD (CRD: aa 240-355) of rfhSP-D and N-terminal of ACE2. The effect of binding of trimeric rfhSP-D to S-protein and dimeric ACE2 on ACE2–S protein interaction was evaluated by further docking the docked complex of S protein and rhfSP-D with ACE2 and ACE2 and rfhSP-D with S protein using Patchdock.

### Expression and Purification of a Recombinant Fragment of Human SP-D Containing Neck and CRD Regions

The rfhSP-D used was expressed and purified from *E. coli* as described previously (18, 19). Briefly, the pUK-D1 plasmid that codes for the 8 Gly-X-Y repeats, neck and CRD regions of human SP-D was transformed into Escherichia coli BL21 (λDE3) pLysS (Invitrogen). The transformed colonies (selected by ampicillin resistance) were grown in Luria-Bertani media supplemented with a final concentration of 100 μg/ml ampicillin and 34 μg/ml chloramphenicol (Sigma-Aldrich) to an OD600 of 0.6. The bacterial culture was then induced to produce the recombinant protein by the addition of 0.5 M isopropyl β-d-1-thiogalactopyranoside (IPTG) (Sigma-Aldrich) and was allowed to grow for a further 3 h. Post incubation, the bacteria were harvested and lysed using lysis buffer (50 mM Tris–HCl pH7.5, 200 mM NaCl, 5 mM EDTA pH 8, 0.1% v/v Triton X-100, 0.1 mM phenyl-methyl-sulfonyl fluoride, 50 μg/ml lysozyme) and sonicated (five cycles, 30 s each). The sonicate was harvested via centrifugation at 12,000 × g for 30 min. This was followed by solubilisation of inclusion bodies in refolding buffer (50 mM Tris–HCl pH 7.5, 100 mM NaCl, 10 mM 2-Mercaptoethanol) containing 8 M urea. and stepwise dialysis of the solubilised fraction against refolding buffer containing 4 M, 2 M, 1 M, and no urea. rfhSP-D was purified from the dialysate by affinity chromatography using a maltose agarose column (5 ml; Sigma-Aldrich). The bound rfhSP-D to the maltose was eluted using elution buffer (50 mM Tris–HCl, pH 7.5, 100 mM NaCl, and 10 mM EDTA) and passed through a PierceTM High-Capacity Endotoxin Removal Resin (Thermofisher) to remove endotoxin. Finally, the endotoxin levels were measured via the QCL-1000 Limulus amoebocyte lysate system (Lonza) and found to be <5 pg/μg of rfhSP-D. The purified rfhSP-D was subjected to western blotting after running on 12% w/v acrylamide SDS-PAGE to assess purity and immunoreactivity (18).

### ELISA

Assays to determine the binding of the S protein or its RBD of SARS-CoV-2 was performed using the SARS-CoV-2 (COVID-19) Inhibitor Screening Kit from Acrobiosystems (EP-105) as per the manufacture’s protocol. Briefly, S protein diluted in Coating Buffer (15 mmol/L sodium carbonate (Na_2_CO_3_), 35 mmol/L sodium hydrogen carbonate (NaHCO_3_), pH 9.6) to a final concentration of 0.3 μg/ml were added to 96 well plates and incubated overnight (~16 h) at 4°C. The uncoated protein was removed by washing the wells with Wash Buffer (PBS with 0.05% (v/v) Tween-20, pH 7.4) three times. The wells were then blocked using the Blocking Buffer (PBS with 0.05% (v/v) Tween-20 and 2% (w/v) bovine serum albumin (BSA), pH 7.4) for 1.5 h at 37°C.

To assess the direct binding of rfhSP-D to S protein, rfhSP-D (20, 10 and 5 μg/ml) were added to the wells. The plate was then incubated for 1 h at 37°C, and any unbound protein was removed by washing the wells three times with the wash buffer. The wells were probed using either polyclonal or monoclonal antibodies against SPD at a dilution of 1:5000 for 1 h at 37°C to detect S protein-rfhSP-D binding. Unbound antibodies were removed by washing three times using the wash buffer. Anti-mouse IgG-Horseradish peroxidase (HRP) (Cat # 31430, Invitrogen), anti-rabbit IgG HRP (Cat # 31466, Invitrogen) or Protein A HRP (Cat # 18-160, Merck) at 1: 5000 dilution was used secondary antibodies by adding them to the respective wells of the appropriate primary antibodies and incubating them for 1 h at 37°C. Following washes with wash buffer three times, the binding was detected using 3,3’,5,5’-Tetramethylbenzidine (TMB) substrate (100 μl/well) (DuoSet ELISA Ancillary Reagent Kit, R&D Systems) as per the manufacturer’s instruction, followed by stopping the reaction using 1M sulphuric acid (100 μl/well) (Cat # Q29307, Thermofisher). The plate was read at 450 nm using a microplate absorbance reader (Synergy H1 multimode plate reader, Biotek). Full-length Surfactant Protein D (FL SP-D) (20 μg/ml) was also used in a similar manner to assess the binding of S protein to FL SP-D. A similar experiment was carried out in parallel using rfhSP-D (20, 10 and 5 μg/ml), supplemented with 10mM EDTA and probed with polyclonal antibodies against SPD (1:5000) to evaluate if the S protein-rfhSP-D binding was calcium-dependent.

The binding of rfhSP-D or FL SP-D to ACE-2 was evaluated using a similar experiment as above. Briefly, FL SP-D (0.1 μg/ml) or rfhSP-D (0.1 μg/ml) were coated in a 96 well plate and probed with decreasing concentration of ACE-2 hACE-2 (0.12, 0.06 and 0.00 μg/ml). The binding was detected using streptavidin tagged with HRP (1:5000) (EP-105, Acrobiosystems), and the colour was developed as described above.

In a separate experiment to assess if rfhSP-D inhibited the interaction between the S protein and biotinylated human Angiotensin-converting enzyme 2 (hACE-2), decreasing concentration of rfhSP-D (5, 1 and 0 μg/ml) preincubated with (hACE-2), were added to wells coated with S protein (0.3 μg/ml) and blocked as described above. The plate was incubated for 1h at 37°C and washed with the wash buffer the times to remove any unbound proteins. The S protein-hACE-2 binding was measured by probing the wells with the HRP tagged Streptavidin antibody (1:5000) for 1h at 37°C. Colour was developed using 3,3’,5,5’-Tetramethylbenzidine (TMB) substrate. The reaction was stopped using 1 M H_2_SO_4_, and the absorbance was read at 450 nm using a microplate absorbance reader. rfhSP-D (5 μg/ml) supplemented with either with 10mM EDTA was used in a similar manner to evaluate if the rfhSP-D mediated inhibition of the interaction between the S protein and biotinylated hACE-2 occurred in a calcium-independent manner. rfhSP-D mediated inhibition of the interaction between the RBD of SARS-CoV-2 S protein and biotinylated hACE-2 was also assessed in a similar manner.

### Vero Cell Infection Assay

Vero cell line (derived from African green monkey epithelial Kidney cells) (ATCC^®^ CCL–81™) (5×10^4^) were cultured for 16 h in each well of a 12 well plate in serum-free medium (MEM Glutamax, containing 1% v/v Penicillin-Streptomycin and 1%v/v sodium pyruvate [Gibco, Thermofisher]). SARS-CoV-2 clinical samples (100 TCID_50_/ well, MOI 0.01) were preincubated with rfhSP-D [0 μg/ml (0 μM), 50 μg/ml (~2.5μM) or 100 μg/ml (~5μM)] in MEM containing 5mM CaCl_2_ for 1h at RT and 1h at 4°C. This pre-treated or untreated virus was added to the cells (Cells + rfhSP-D + Virus). After 1h incubation at 37°C, 5% CO_2_, the medium was removed, and cells were washed with PBS to remove any unbound CoVs. Infection medium (MEM+0.3% BSA) was added to the cells and incubated for 24 h to assess replication. The cells were then harvested by scraping with a sterile disposable cell scraper and centrifuged at 1500 x g for 5 minutes. Total RNA was extracted using the Perkin Elmer automated extractor and subjected to Real-time RT-PCR for SARS-CoV-2 using Pathodetect kits from MyLabs, as per manufacture’s protocol. For the replication analysis of SARS-CoV-2, Ct value for SARS-CoV-2 RNA dependent RNA polymerase (RdRp) gene was used for analysis. Cells incubated with rfhSP-D, without virus was used protein control (Cells + rfhSP-D) and cells incubated with BSA (100μg/ml), and the virus was used as non-specific protein control (Cells + Virus). Sterile PBS with the virus was used as negative control.

The effect of rfhSP-D on viral infection was assessed by culturing Vero cells (5×10^5^) in a 12 well plate in serum-free MEM. SARS-CoV-2 clinical samples (500 TCID_50_/ well, MOI 0.05) were treated with rfhSP-D and added to the cells as described above. However, after the addition of the infection medium, the cells were incubated only for 2h, after which they were harvested, and Real-time RT-PCR was performed using the same controls and parameters described above.

### Statistical Analysis

Graphs were generated using GraphPad Prism 8.0 software, and the statistical analysis was performed using a two-way ANOVA test. Significant values were considered based on *p < 0.1, **p < 0.05, ***p < 0.01, and ****p < 0.001 between treated and untreated conditions. Error bars show the SD or SEM, as indicated in the figure legends.

## Results

### rfhSP-D interacts with the Spike protein of SARS-CoV-2 and human ACE-2 in *silico*

S protein is known to interact via the receptor binding motif (RBM:455-508) in the receptor binding domain (RBD: aa 319-527) with virus binding hotspot residues comprising of Lys31, Glu35 and Lys353 of dimeric hACE2 (14–16). The structure of hACE2 receptor, co-crystallized with Spike S protein of SARS-CoV-2, is available in RCSB (pdb id: 6VW1). The receptor (ACE2) and ligand (Spike S) were separated and docked to validate the docking protocol. The redocked complex of ACE2 and S protein had root mean square deviation (RMSD) of 7.9 Å. The close agreement between the docked and crystal structures validated the docking protocol used in the study.

In case of docked solutions for S protein and rfhSP-D **(Supplementary Figure 1)**, the third ranked docked pose with binding energy of −20.63 kcal/mol exhibited rfhSP-D interactions with RBM residues Tyr449, Gln493 Gln498, implying that rfhSP-D could bind to Spike protein in a manner that can inhibit ACE2-S protein interaction (**Table 2**; **Figure 1**). To ascertain this hypothesis, the complex of S protein with rfhSP-D was docked to ACE2. S protein and rfhSP-D bound to ACE2 via common interacting residues.

**Figure 1:**
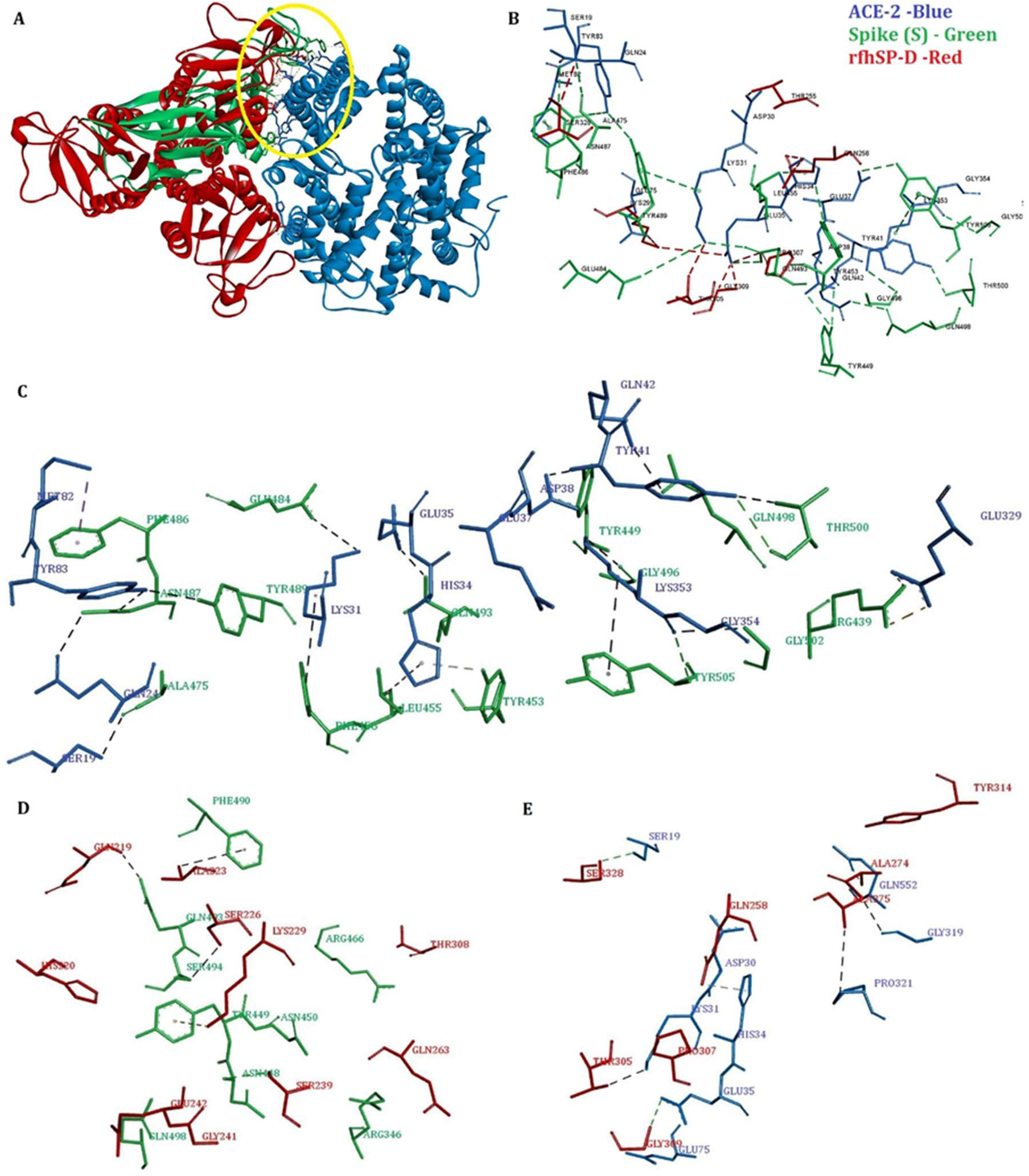
Tripartite interaction between S protein (Green), rfhSP-D (Red) and ACE-2 (Blue) [A, B (zoomed view)]. ACE-2 residues, Ser19, Lys31, Glu35 and His34, interact with both S protein and rfhSP-D. The interactions between S protein and ACE-2 are deduced from the crystal structure (PDB ID: 6VW1) and between rfhSP-D, and ACE-2 protein are based on docked complexes. Individual intermolecular interactions between **(C)** S protein (Green) and ACE-2 (Blue); **(D)** S protein (Green) and rfhSP-D (Red) and **(E)** rfhSP-D (Red) and ACE-2 (Blue). The S protein residues, Tyr449, Gln493 and Gln498, participate in intermolecular interactions with both ACE-2 and rfhSP-D.

**Table 2:** Results of docking of S protein, ACE-2, and rfhSP-D

| S. No | Receptor | Ligand | Binding energy (kcal/mol) | Interactions |  |
| --- | --- | --- | --- | --- | --- |
| | | | | Receptor<sup>\$</sup> | Ligand* |
| 1 | ACE-2 | S protein | Crystal structure | **Ser19** | Ala475 |
|  |  |  |  | Gln24 | Asn487 |
|  |  |  |  | **Lys31** | Phe456, Glu484, Tyr489, **Gln493** |
|  |  |  |  | **His34** | Leu455, Tyr453 |
|  |  |  |  | **Glu35** | **Gln493** |
|  |  |  |  | Glu37 | Tyr505 |
|  |  |  |  | Asp38 | **Tyr449** |
|  |  |  |  | Tyr41 | Thr500 |
|  |  |  |  | Gln42 | **Gln498** |
|  |  |  |  | Met82 | Phe486<sup>@</sup> |
|  |  |  |  | Tyr83 | Gly496, Asn487, Tyr489 |
|  |  |  |  | Glu329 | Arg439 |
|  |  |  |  | Lys353 | Tyr505, Gly502 |
|  |  |  |  | Gly354 | Tyr505 |
| 2 | rfhSP -D | S protein (Open) | -20.63 | Gln219 | **Gln493** |
|  |  |  |  | His220 | **Tyr449** |
|  |  |  |  | Ala223 | **Gln493**, Phe490 |
|  |  |  |  | Ser226 | Ser494 |
|  |  |  |  | Lys229 | **Tyr449** |
|  |  |  |  | Ser239 | Asn450 |
|  |  |  |  | Gly241 | Asn448, **Gln498** |
|  |  |  |  | Glu242 | **Gln498** |
|  |  |  |  | Gln263 | Arg346 |
|  |  |  |  | Thr308 | Arg466 |
| 3 | ACE-2 | rfhSP-D | -24.30 | **Ser19** | Ser328 |
|  |  |  |  | Asp30 | Thr255 |
|  |  |  |  | **Lys31** | Thr305 |
|  |  |  |  | **His34** | Gln258 |
|  |  |  |  | **Glu35** | Pro307, Gly309 |
|  |  |  |  | Glu75 | Lys299 |
|  |  |  |  | Gly319 | Ala275 |
|  |  |  |  | Pro321 | Ala275 |
|  |  |  |  | Gln552 | Ala274, Tyr314 |

\*The S protein residues in bold are predicted to be part of the common binding site for ACE2 and rfhSP-D.
<sup>\$</sup>The ACE2 residues in bold interact with both S protein and rfhSP-D (docked structure).
<sup>@</sup>The structural coordinates of Phe486 is missing in the open conformation S protein (PDB ID: 6VYB).

The top ranked docked structure of ACE2 and rfhSP-D had binding energy of −24.30 kcal/mol. In this pose, rfhSP-D interacted with the virus-binding hotspot residues Ser19, Lys31, His34 and Glu35 of ACE2, implying that rfhSP-D could bind to ACE2 in a manner that can inhibit ACE2-S protein interaction **(Table 2, Figure 1)**. To corroborate this postulation, the complex of ACE2 with rfhSP-D was docked to Spike S. Top ranked pose of ACE2-rfhSP-D complex docked with open S protein had binding energy of −33.01 kcal/mol and several common interactions between rfhSP-D and ACE2 with S protein **(Supplementary Figure 1**). The docking experiments led us to infer that rfhSP-D could bind to both ACE2 and Spike S and prevent ACE2-S protein interaction.

### rfhSP-D binds to the immobilised S protein of the SARS-CoV-2 as well as hACE-2

The possible binding between rfhSP-D and S protein hinted by the docking analysis was confirmed *in vitro* via an indirect ELISA. rfhSP-D was found to bind the immobilised S protein in a dose-dependent manner **(Figure 2a)**. However, a significant difference in the absorbance was observed based on the specificity of the primary antibody used. S protein-rfhSP-D binding that was probed with the polyclonal antibody against SP-D reported a significantly higher absorbance when compared to the wells that were probed with a monoclonal antibody directed against the CRD of SP-D. This difference suggests involvement of CRD of rfhSP-D with the spike protein and therefore, the CRD was not available for interaction with the monoclonal antibody. S protein was also found to bind to the FL SP-D. The treatment of rfhSP-D with 10mM EDTA did not significantly alter the binding of rfhSP-D to S protein **(Figure 2b)**. Hence, rfhSP-D binds to the S protein in a dose-dependent but a calcium-independent manner. A similar parallel experiment revealed that rfhSP-D bound ACE2 in a dose-dependent manner **(Figure 2c)**.

**Figure 2:**
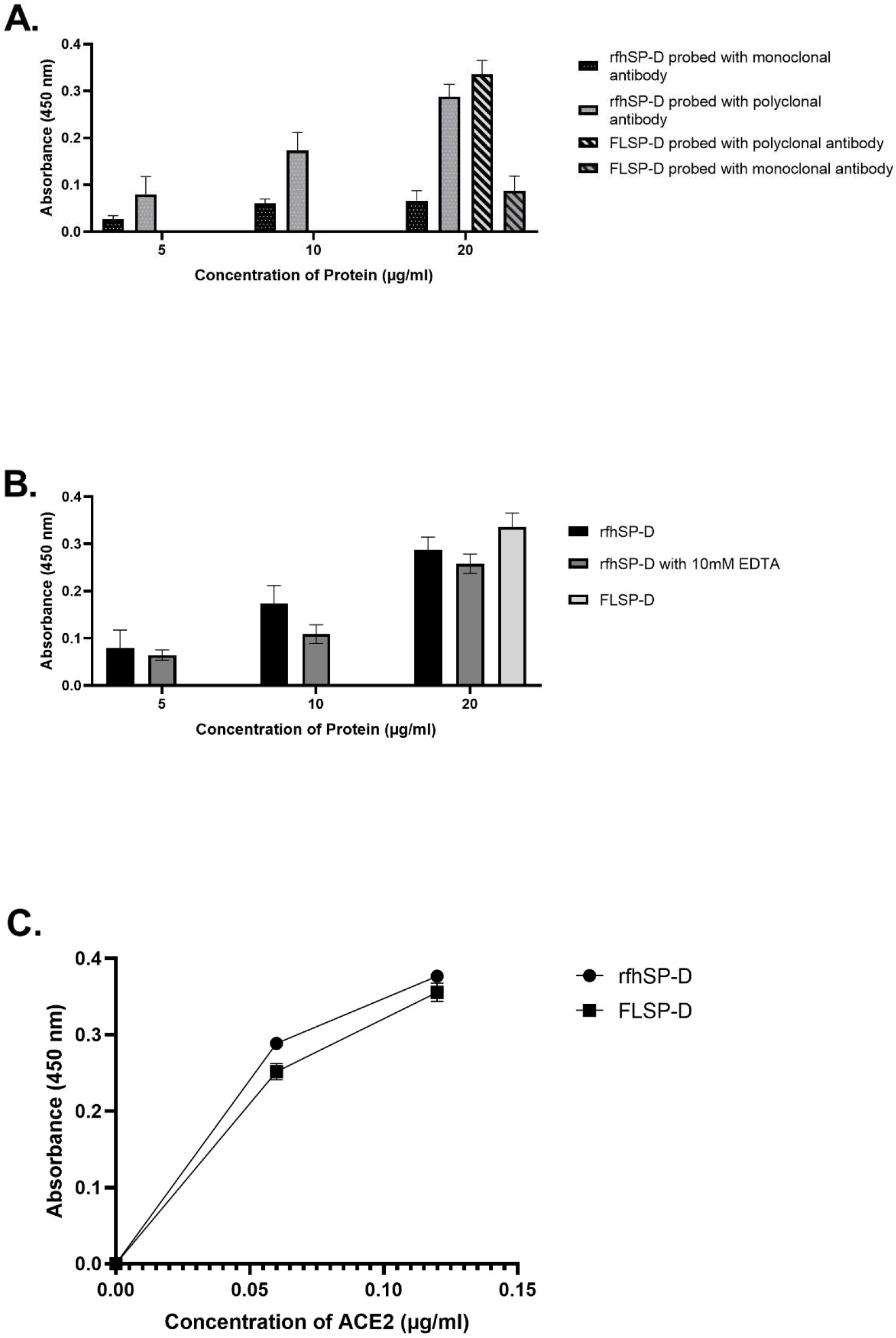
rfhSP-D binds to the immobilised Spike protein (S protein) of the SARS-CoV-2; immobilised rfhSP-D binds to hACE-2 in a dose-dependent but calcium-independent manner. ELISA showing binding of rfhSP-D to the immobilised S protein in a dose-dependent manner. Microtiter wells were coated with 0.3 μg/ml of S protein. rfhSP-D (20, 10 and 5 μg/ml) were added to the wells. Full-length Surfactant Protein D (FL SP-D) (20 μg/ml) was also used in a similar manner. S protein-SP-D binding was detected with either polyclonal or monoclonal **(A)** antibodies against SP-D. To assess the effect of calcium in the rfhSP-D-S protein interaction, rfhSP-D either with/without 10mM EDTA was used in a similar manner and probed with polyclonal antibodies against SP-D **(B)**. The binding of immobilised rfhSP-D to hACE-2 **(C)** was assessed by coating microtiter wells with 0.1 μg/ml of FL SP-D or rfhSP-D. Decreasing concentration of hACE-2 (0.12, 0.06 and 0.00 μg/ml) was added to the wells. The SP-D-hACE-2 binding was detected with Streptavidin-HRP. The background was subtracted from all data points. The data were expressed as the mean of triplicates ± SD.

### rfhSP-D inhibits the interaction of S protein and its RBD with biotinylated hACE-2 in a calcium-independent manner

Since rfhSP-D was found to bind to the S protein and ACE-2, and as both rfhSP-D and ACE-2 were predicted to share the same binding site on S protein, rfhSP-D mediated inhibition of the interaction between the RBD of S protein of SARS-CoV-2 and ACE-2 was assessed using a colorimetric ELISA.

The wells were coated with either the S protein or its RBD domain that was preincubated with rfhSP-D followed by biotinylated hACE-2. The functionality and the range of the assay were initially assessed by verifying if the assay could detect the binding of hACE-2 at a concentration of 0.12 μg/ml and 0.06 μg/ml. The binding occurred in a dose-dependent manner, confirming that the assay can detect binding between S protein or its RBD domain with hACE-2 at a concentration as low as 60 ng/ml **(Supplementary Figure 2)**. A decrease in binding between S protein and hACE-2 was observed as the concentration of rfhSP-D increased **(Figure 3 and Figure 4)**. Approximately 50% decrease in S protein-hACE-2 binding was observed as rfhSP-D concentration increased 5-fold **(Figure 3a)**. A similar result was observed between the binding of the RBD of S protein and hACE-2. An 8-fold increase in the concentration of rfhSP-D was found to decrease RBD-hACE-2 interaction by ~25% **(Figure 4a)**. No significant difference was observed between the samples with 10mM EDTA and without EDTA in terms of rfhSP-D mediated S protein/RBD-hACE-2 binding **(Figure 3b; Figure 4b)**. Hence, rfhSP-D mediated inhibition of the interaction between the RBD of S protein or the S protein itself with biotinylated hACE-2 occurred in a calcium-independent manner.

**Figure 3:**
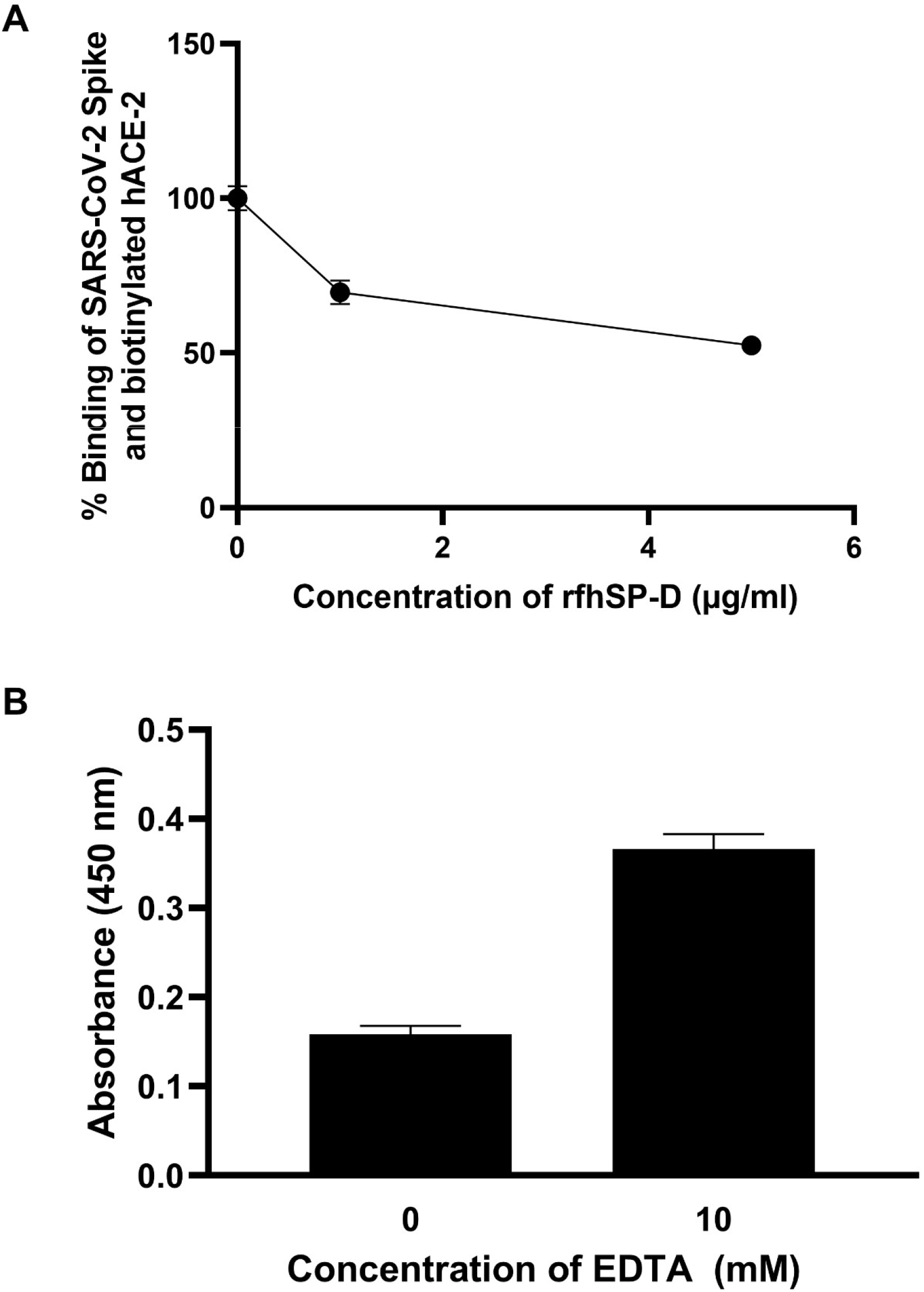
rfhSP-D inhibits the interaction between Spike of SARS-CoV-2 and biotinylated hACE-2 in a calcium-independent manner. Microtiter wells were coated with 0.3 μg/ml of S protein. rfhSP-D (5, 1 and 0 μg/ml) **(A)** pre-incubated with biotinylated human Angiotensin-converting enzyme 2 (hACE-2) was added to the wells. To assess the effect of calcium in the rfhSP-D-mediated inhibition of S protein-hACE-2 interaction **(B)**, 5 μg/ml of rfhSP-D with/without 10 mM EDTA. S protein-hACE-2 binding was detected with Streptavidin-HRP. Background was subtracted from all data points. The data were normalised with 100% S protein: hACE-2 binding being defined as the mean of the absorbance recorded from the control sample (0 μg/ml of rfhSP-D). The data were presented as the mean of the normalised triplicates ± SEM for inset A and B. The data were presented as the mean of the triplicates ± SEM for inset C. Significance was determined using the two-way ANOVA (n = 3); no significant difference was observed between the samples with 10mM EDTA and without EDTA in terms of rfhSP-D-mediated S protein:hACE-2 binding.

**Figure 4:**
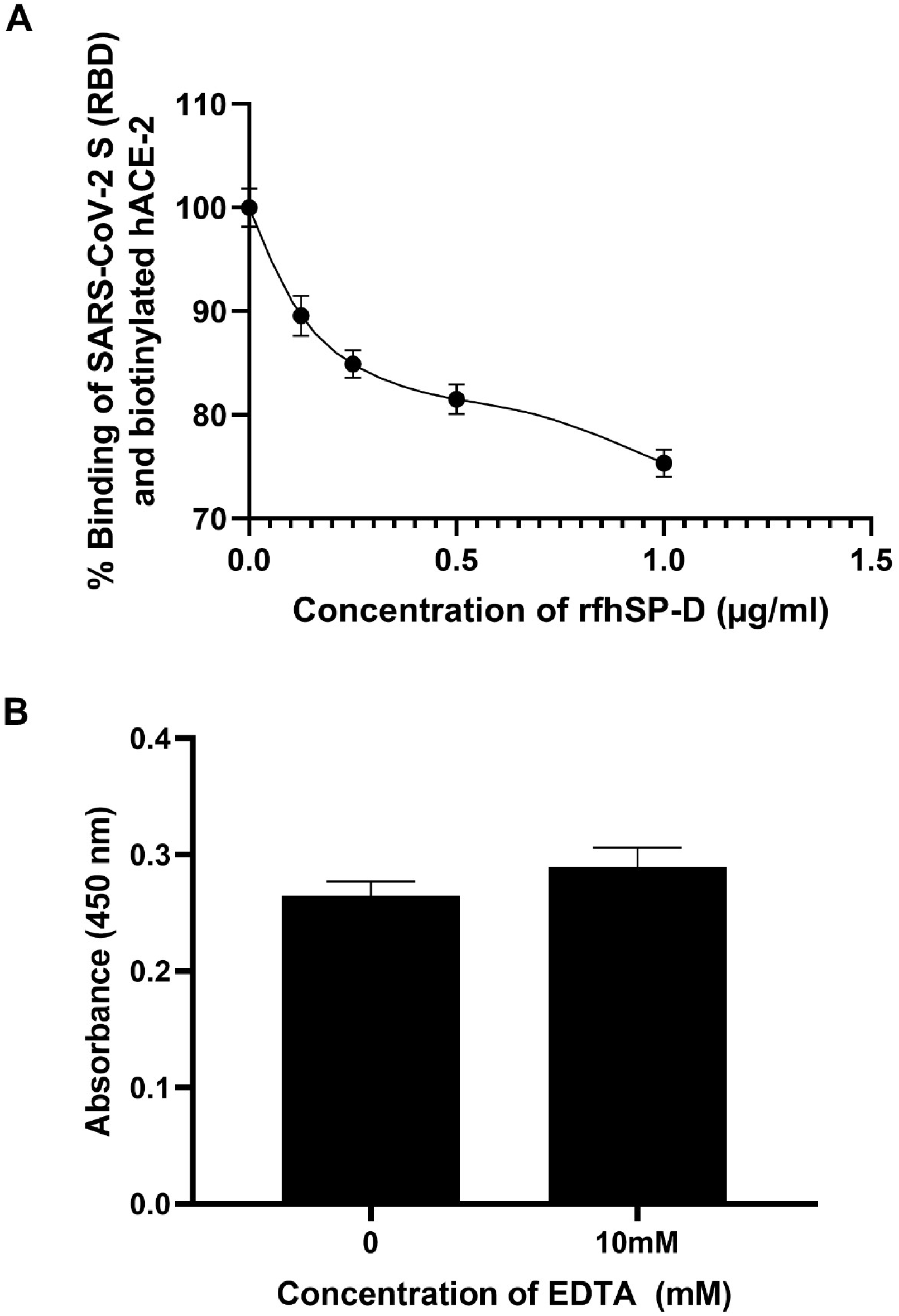
rfhSP-D inhibits the interaction between RBD of Spike protein of SARS-CoV-2 and biotinylated hACE-2 in a calcium-independent manner. Microtiter wells were coated with 0.1 μg/ml of S protein RBD. Decreasing concentration of rfhSP-D (1, 0.5, 0.25, 0.125, and 0 μg/ml **(A)** co-incubated with biotinylated human Angiotensin-converting enzyme 2 (hACE-2) was added to the wells. To assess the effect of calcium in the rfhSP-D-mediated inhibition of S protein RBD: hACE-2 interaction **(B)**, 5 μg/ml of rfhSP-D with/without 10mM EDTA was used. S protein RBD-hACE-2 binding was detected with Streptavidin-HRP. Background was subtracted from all data points. The data obtained were normalised with 100% S protein RBD-hACE-2 binding being defined as the mean of the absorbance recorded from the control sample (0 μg/ml of rfhSP-D). The data were presented as the mean of the normalised triplicates ± SEM for inset A and B. The data were presented as the mean of the triplicates ± SEM for inset C. Significance was determined using the two-way ANOVA (n = 3) and no significant difference was observed between the samples with 10mM EDTA and without EDTA in terms of rfhSP-D mediated S protein RBD: hACE-2 binding.

### rfhSP-D treatment inhibits SARS-CoV-2 infection and replication

As rfhSP-D is known to induce apoptosis in cancer and immortalised cells (18, 20–22), the effect of rfhSP-D on Vero cells was assessed using MTT assay. rfhSP-D treatment had no significant effect on the viability of Vero cells (**Supplementary figure 3**). At the outset, the TCID_50_ values of the clinical samples were obtained by evaluating the cytopathic effects using MTT assay. As expected, when Vero cells were challenged with 100 TCID_50_, or 50 TCID_50_ of viral samples from SARS-CoV-2 clinical samples, a 50% or 25% reduction in cell viability was observed, respectively, compared to the viability of uninfected Vero cells, confirming the assayed TCID_50_ values (**Figure 5)**. The control samples showed no significant difference in the cell viability than the uninfected Vero cells when the control sample volumes equivalent to 100 TCID_50_ and 50 TCID_50_ of the matched clinical cases were used.

**Figure 5:**
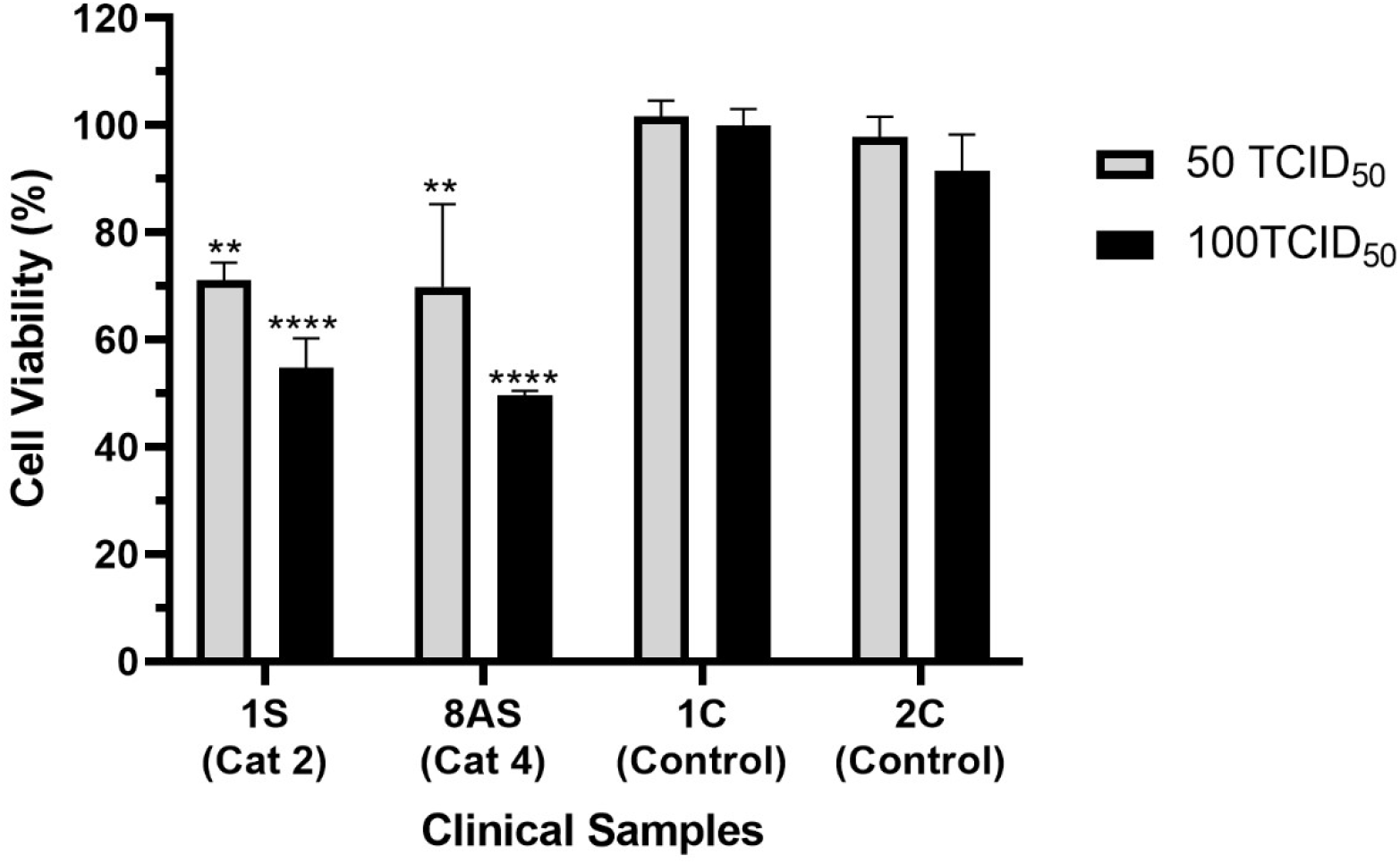
Determination of TCID_50_ value of the clinical samples in Vero cells using MTT assay. Vero cells (5 x 10^4^/well) were seeded in complete MEM in 96-well culture plates and grown overnight at 37°C with 5% CO_2_. Swab samples of 15 confirmed cases of COVID-19 (n=5 of Cat2, Cat6 and Asymptomatic) and 15 controls (at different dilutions/well) were added to the cells and incubated for 1h. The supernatants were removed, and the wells were washed twice with sterile PBS. Fresh complete MEM was added to the wells, and the cells were incubated for 96 h. Viability of the cells was evaluated using MTT assay. MTT (0.5 mg/ml) containing medium was added to the wells for 4h. The supernatants were removed, and cells were lysed using DMSO. Absorbance was measured at 590nm. The data obtained were normalised with 100% cell viability being defined as the mean of the absorbance recorded from the control sample (0 TCID_50_/well) and TCID_50_ units were evaluated in each sample. The same assay was used to validate the cytopathic effects of 100TCID_50_ and 50TCID_50_ units of the samples. The representative data for cases (n=2) and controls (n=2) are presented as the mean of the normalised triplicates ± SEM Significance was determined using the two-way ANOVA (n = 3) test (**p < 0.01, and \*\*\*\**p* < 0.0001).

The effect of rfhSP-D on the replication of SARS-CoV-2 (100 TCID_50_/well; MOI 0.01) in Vero cells was evaluated by measuring the levels of the RdRp gene of SARS-CoV-2 by RT-PCR 24h post-infection. Pre-treatment of the positive samples (n=15), comprising of SARS-CoV-2 with rfhSP-D, led to a reduction in RdRp levels in a dose-dependent manner **(Figure 6; Table S1)**. The pre-treatment of samples from all categories of SARS-CoV-2 positive cases [as representatives, the figure 6 shows the data for 1S (Cat 2) and 3S (Cat 6)] with 2.5μM rfhSP-D led to ~4.5-fold reduction (−4.5 log2) of RdRp transcript compared to the untreated positive sample challenged Vero cells. There was no significant difference in the Ct values of RdRp gene from the untreated and control sample treated Vero cells. Similarly, pre-treatment with 5 μM rfhSP-D resulted in ~5.5-fold reduction (−5.5 log2) of RdRp mRNA expression. Remdesivir, one of the anti-viral drugs proposed for COVID-19, which functions by inhibiting viral RNA synthesis, was found to inhibit SARS-CoV-2 replication by ~4-fold (−4 log2). Hence, rfhSP-D blocked SARS-CoV-2 infection, in addition to inhibiting the replication of SARS-CoV-2 significantly better than Remdesivir at both tested concentrations (2.5 μM and 5 μM rfhSP-D).

**Figure 6:**
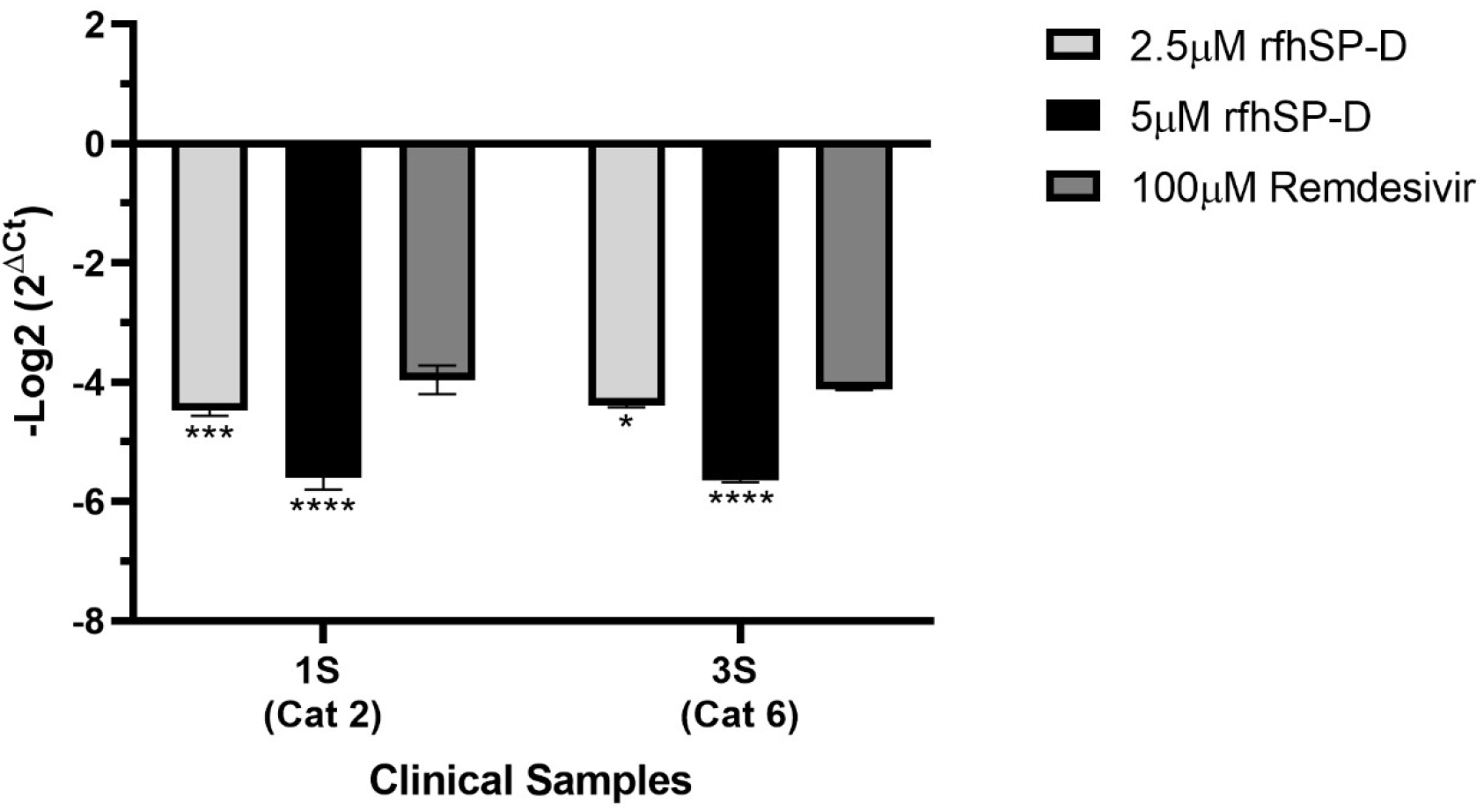
rfhSP-D pre-treatment of SARS-CoV-2 significantly inhibited its replication. Vero cells (5 x 10^4^/well) were seeded in complete MEM in 96-well culture plates and grown overnight at 37°C under 5% CO_2_. Cells were washed with sterile PBS twice. SARS-CoV-2 clinical samples (100TCID_50_/ well; MOI 0.01) were preincubated with rfhSP-D [0 μg/ml (0 μM), 50 μg/ml (~2.5μM) or 100 μg/ml (~5μM)] in MEM containing 5mM CaCl_2_ for 1h at RT. The pre-treated or untreated virus in the sample was added to the cells and incubated for 1h at 37°C under 5% CO_2_. The wells were washed with PBS twice, and infection medium (MEM+0.3% BSA) was added to the cells and incubated for 24h at 37°C. The supernatants were collected, RNA was extracted by Perkin Elmer automated extractor, and subjected to qRT-PCR for SARS-CoV-2. For control samples, the volume of the sample taken was equivalent to the volume of the case sample (100 TCID_50_) where no RdRp expression was detected. The relative expression of RdRp was calculated using rfhSP-D untreated cells (0 μM rfhSP-D), infected with respective samples as the calibrator. Data of representative cases (n=2) is presented as the mean of triplicates (n=3). Error bars represent ± SEM. Significance (compared to 100μM Remdesivir) was determined using the two-way ANOVA test (\**p* < 0.05, \*\*\**p* < 0.01, and \*\*\*\**p* < 0.0001).

As rfhSP-D was found to interact with S protein and ACE-2, proteins that play an integral role in viral host cell recognition and entry, the role of rfhSP-D in viral infectivity was assessed in a similar manner to replication. Vero cells infected with SARS-CoV-2 positive samples (500TCID_50_/well, MOI 0.05) showed a rfhSP-D dose-dependent decrease in the expression levels of the RdRp gene, 2h post-infection **(Figure 7; Table S2)**. Clinical samples from all the categories of SARS-CoV-2 patients [As representatives, figure 7 shows the data from 2S (Cat 6) and 9AS (Cat 5a)] showed ~ 1.25-fold reduction (−1.25 log2) or ~ 2-fold reduction (−2 log2) in RdRp gene expression with the samples pre-treated with either 2.5μM or 5 μM respectively of rfhSP-D. Remdesivir was used as a control (Remdesivir does not inhibit SARS-CoV-2 infection). Thus, pre-treatment of SARS-CoV-2 in the clinical sample with rfhSP-D appears to make S protein unavailable to interact with the ACE-2 receptor on the host cell, thus, reducing the infectivity of the virus and subsequent viral replication in a dose-dependent manner.

**Figure 7:**
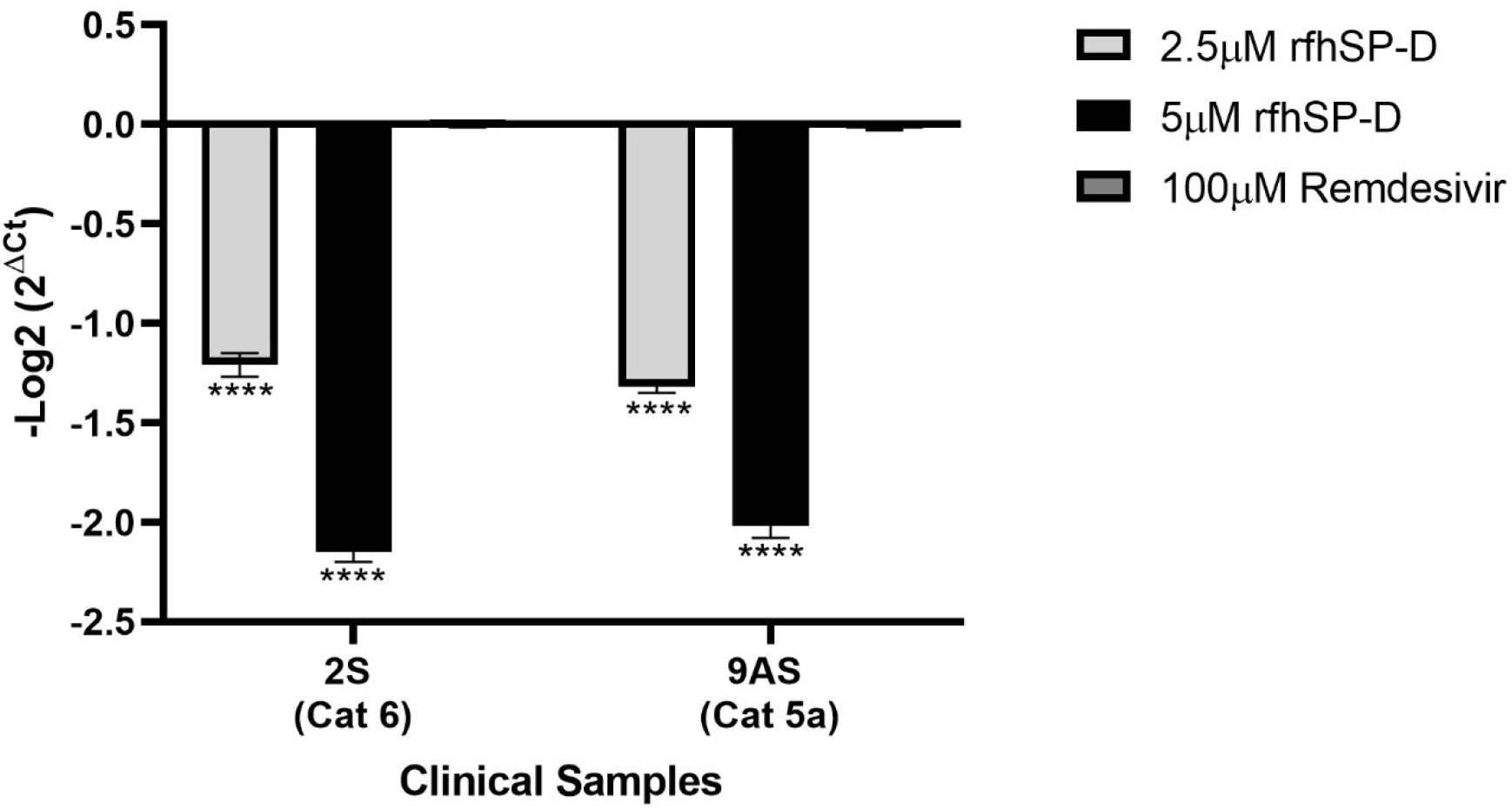
rfhSP-D pre-treatment of SARS-CoV-2 significantly inhibited its infectivity. Vero cells (5 x 10^5^/well) were seeded in complete MEM in 12-well culture plates and grown overnight at 37°C under 5% CO_2_. Cells were washed with sterile PBS twice. SARS-CoV-2 clinical samples (500TCID_50_/well, MOI 0.05) were preincubated with rfhSP-D [0 μg/ml (0 μM), 50 μg/ml (~2.5μM) or 100 μg/ml (~5μM)] in MEM containing 5mM CaCl_2_ for 1h at RT and 1h at 4°C. This pre-treated or untreated virus containing sample was added to the cells and incubated for 1h at 37°C under 5% CO_2_. The wells were washed with PBS twice, and infection medium (MEM+0.3% BSA) was added to the cells and incubated for 2h at 37°C under 5% CO_2_. The cells were scraped, and the media containing scraped cells were collected. RNA was extracted and subjected to RT-PCR for SARS-CoV-2. For control samples, the volume of the sample taken was equivalent to the volume of the case sample (500 TCID_50_); no RdRp expression was detected the relative expression of RdRp was calculated by using rfhSP-D untreated cells (0 μM rfhSP-D), infected with respective samples as the calibrator. Data for representative cases (n=2) is presented as the mean of triplicates (n=3). Error bars represent ± SEM. Significance [compared to control sample (Cells + Virus)] was determined using the two-way ANOVA test (\*\*\*\**p* < 0.0001).

## Discussion

The present study explored the likely protective effect of a recombinant fragment of human lung surfactant protein D, rfhSP-D, against SARS-CoV-2. As predicted by the docking study, rfhSP-D interacted with the spike protein of SARS-CoV-2, its receptor binding domain (RBD) as well as ACE-2. Importantly, these interactions may have contributed to significant inhibition of infectivity and replication of SARS-CoV-2 virus present in the clinical samples derived from asymptomatic, symptomatic and severe patients of COVID-19.

One of the first steps of the SARS-CoV-2 infection is the binding of the S protein to the host cell via, ACE-2 receptor (23). S1 protein is known to interact with ACE-2 receptor via the receptor-binding motif (RBM:455-508) in the receptor-binding domain (RBD: aa 319-527) with virus binding hotspot residues comprising of Lys31, Glu35 and Lys353 of dimeric ACE2 (24–26). Since SP-D interaction with spike protein of SARS-CoV has been reported, which shares ~74% homology with the RBD of SARS-CoV-2 (12) and rfhSP-D is known to bind to viral surface proteins such as haemagglutinin and neuraminidase of influenza A virus, gp120 of human immunodeficiency virus 1 (7, 27), and S protein of SARS-CoV (12), the possibility of rfhSP-D binding to the S protein of SARS-CoV-2 was examined.

*In-silico* interaction of rfhSP-D with RBD of Spike protein of SARS-CoV-2 revealed that Tyr449, Gln493 and Gln498 of RBD overlapped with the residues that are essential for the binding of S protein to the target protein ACE-2. The binding of S protein to rfhSP-D or FL SP-D was confirmed using an indirect ELISA. A comparatively lower absorbance with the monoclonal antibodies raised against the CRD region of human SP-D than the polyclonal antibodies could be attributed to the fact that the binding between rfhSP-D and S protein occurred through the CRD region of rfhSP-D and FL SP-D. Further, calcium independence suggested an involvement of protein-protein interaction. A significant inhibition of the S protein-ACE-2 interaction in presence of rfhSP-D suggested that rfhSP-D could interfere with the binding of the SARS-CoV-2 to the host cell, an essential step for the infection to occur.

Clinical samples of SARS-CoV-2 were used to assess if rfhSP-D modulated the infectivity and replication of the virus (isolation of the virus in the laboratory conditions may introduce alterations) *in vitro*. For assessing replication, qRT-PCR of the RdRp gene, which is essential for the replication of viral RNA was measured. Remdesivir was used as a positive control for replication inhibition. Remdesivir, an adenosine analogue, functions by incorporating itself into nascent viral RNA chains which results in premature termination, thereby effectively inhibiting viral RNA synthesis (28). Downregulated RdRp expression in the Remdesivir treated samples clearly validated the platform for evaluating viral replication using clinical samples. A dose-dependent reduction of the RdRp mRNA expression in Vero cells, challenged with rfhSP-D-pre-treated SARS-CoV-2 positive clinical samples at a higher fold change than Remdesivir, suggested a highly potent anti-SARS-CoV-2 activity mediated by rfhSP-D.

Replication kinetic studies involving Vero cells infected with SARS-CoV-2 have demonstrated a significant synthesis of viral RNA at ≥ 6 h post-infection (29). As such, any viral RNA detected 1-2 h post-infection could be considered to have come from the infecting viral particles and not from subsequent viral RNA synthesis or viral replication. Hence, to confirm if rfhSP-D played a role in inhibiting SARS-CoV-2 infection, Vero cells were infected with SARS-CoV-2 clinical samples at a high concentration (500TCID_50_; MOI 0.05) for 2h. In accordance with the previous reports, no significant effect of Remdesivir on the Ct values of Vero cells challenged with clinical samples validated the assay format (30). Reduced RdRp transcripts in presence of rfhSP-D demonstrated the ability of rfhSP-D to act as an entry inhibitor against SARS-CoV-2. These results suggest that rfhSP-D is a potential candidate to be used as an S protein-based inhibitor against SARS-CoV-2 infections. With established safety *in vivo* and therapeutic efficacy against several respiratory pathogens, rfhSP-D will effectively combat the nosocomial co-infections in COVID-19 patients.

There is dysregulated pro-inflammatory cytokine response without protective IFNs in response to SARS-CoV-2 mediated lung tissue damage leading to Acute Respiratory Distress Syndrome (ARDS). The levels of SP-D were significantly altered in bronchoalveolar lavage of patients of ARDS and were strong predictors of poor prognosis (31, 32). Persistent complement activation leads to microangiopathy leading to hypoxia in vital organs. The current therapeutic strategy comprises of an antiviral like Remdesivir and immunosuppressants such as corticosteroids. Importantly, there is a need to rapidly clear cell debris or Damage Associated Molecular Patterns (DAMPs) and polarise protective immune response towards a protective one and regulate the complement activation. The rfhSP-D is capable of dampening the ‘Cytokine storm’ by rapid clearance of the virus infected cells and strengthening the lung capacity by restoring homeostasis (33).

rfhSP-D has been previously shown to inhibit HIV-1 entry as well successfully thwart the cytokine storm in an ex vivo model of human vaginal tissue (27). SP-D has a compelling role in correcting lung pathophysiology and injury (34). It is possible that SP-D functions as an opsonin after binding to the S protein and helps in viral clearance. As a complement- and antibody-independent neutralisation agent against SARS-CoV-2, rfhSP-D may be a viable alternative as an inhalation formulation to control COVID-19 infection in immunocompromised/deficient people and other populations where vaccination against the virus would not be a viable option. These promising results warrant further studies in COVID-19 animal models, such as mice humanised with human ACE2 and Syrian hamsters (*Mesocricetus auratus*), to better understand the impact of rfhSP-D in the microenvironment of the respiratory system (35).

## Ethics Statement

The project was approved by the institutional ethics committee of Institute of Liver and Biliary Sciences, Delhi (IEC/2020/80/MA04) on 20^th^ July 2020. The committee waived off the written informed consent in due consideration of the request as these samples were stored in the facility and anonymised aliquots of the samples were provided for the study.

## Acknowledgments

Authors acknowledge the support of Dr. Smita Mahale, Director, ICMR-NIRRH; Dr. Shiv K. Sarin Director, ILBS. The study was reviewed and recommended by ICMR Expert Review Committee, and authors acknowledge the support of Dr. Nivedita Gupta and Dr. Raman Gangakhedkar of ICMR-ECD. The study was reviewed by BIRAC-PACE expert panel.

## Supplementary Figures

**Supplementary Figure S1:**
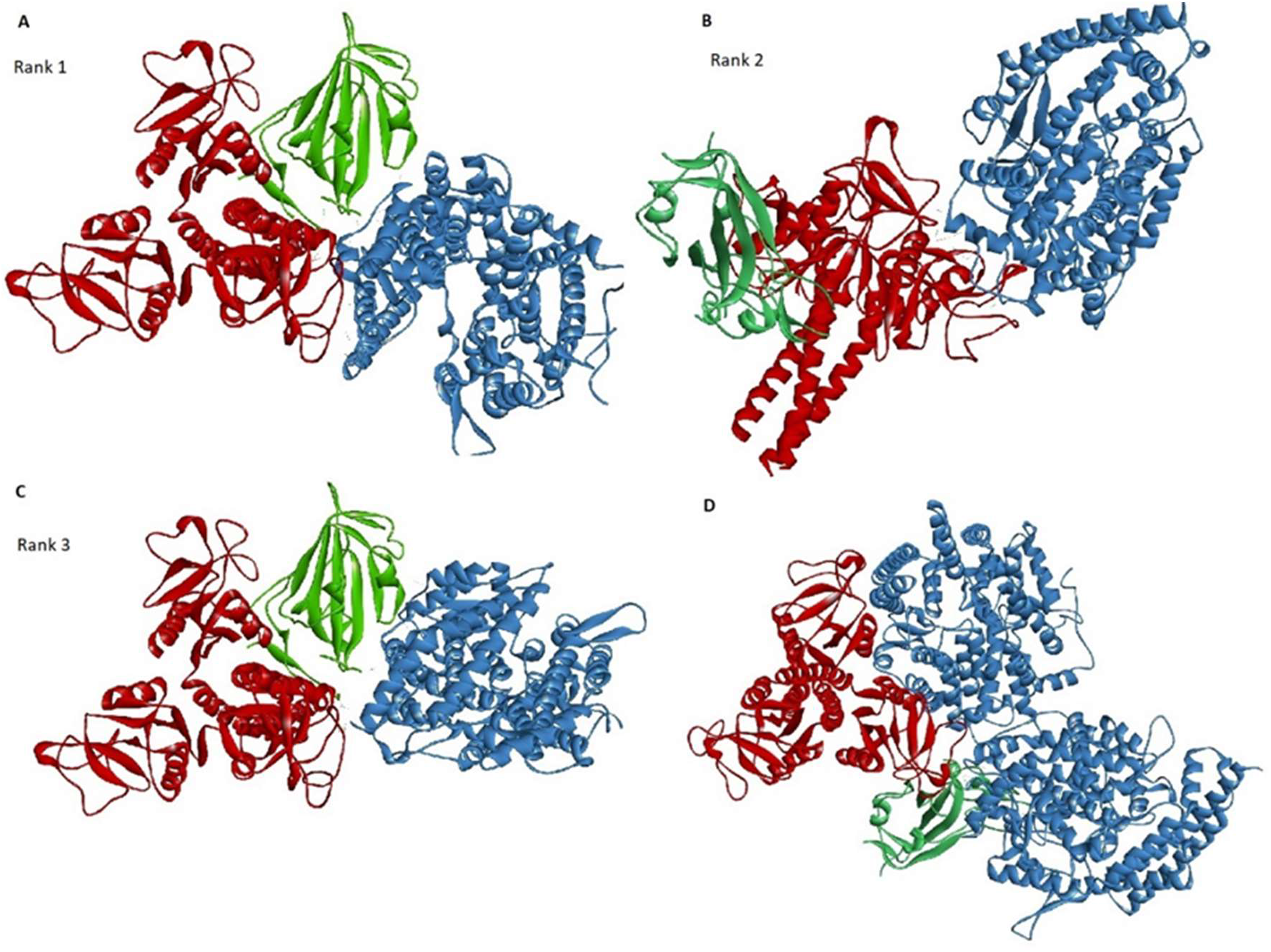
Docked poses of S protein (Green) and rfhSP-D (Red) complex with ACE2 (Blue) (A-C) and ACE2 and rfhSP-D complex with S protein (D).

**Supplementary Figure S2:**
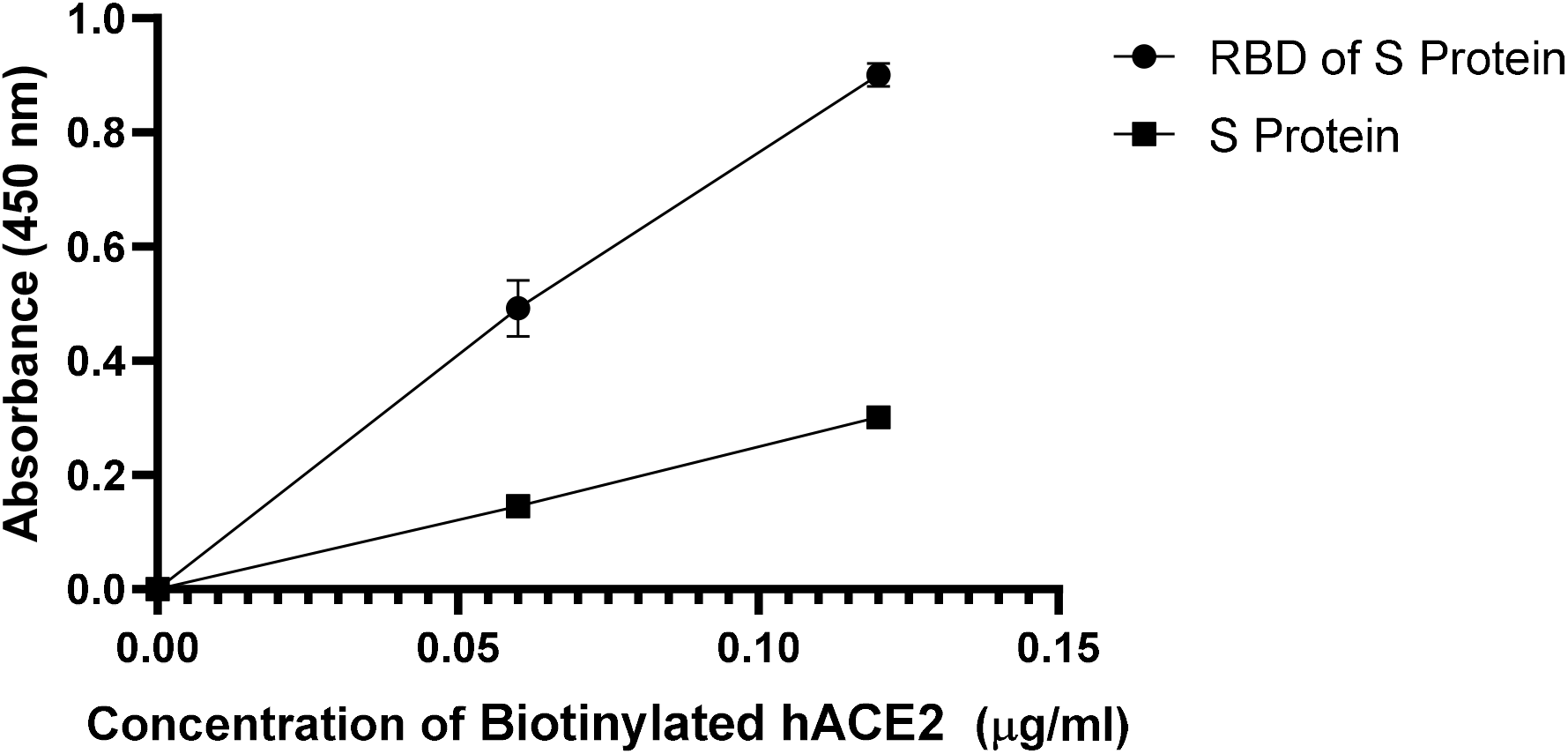
ELISA showing binding of Biotinylated human Angiotensin-converting enzyme 2 (hACE-2) to immobilised SARS-CoV-2 Spike protein (S protein) and the Ribosome Binding Domain (RBD) of the S protein in a linear range. Microtiter wells were coated with 0.3 μg/ml of S protein, or 0.1 μg/ml of RBD of S protein. Decreasing concentration of hACE-2 (0.12, 0.06 and 0.00 μg/ml) were added to the wells. S protein or RBD: hACE-2 binding was detected with Streptavidin-HRP. Background was subtracted from all data points. The data were expressed as the mean of triplicates ± SD.

**Supplementary Figure S3:**
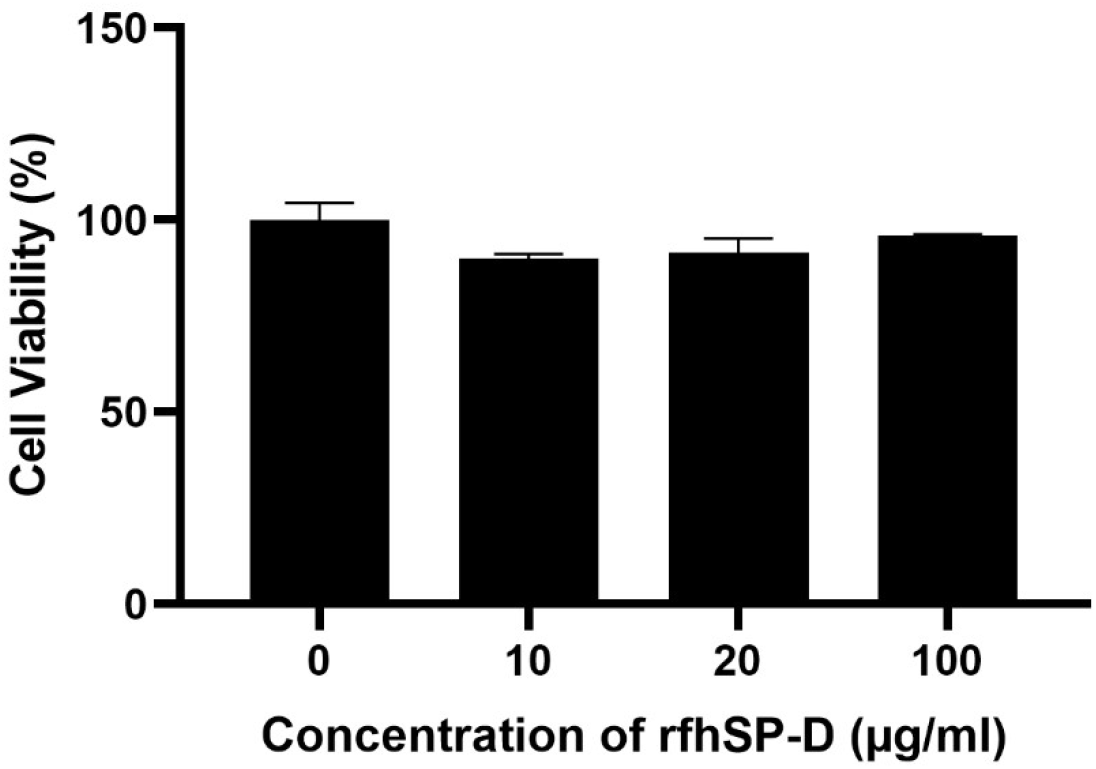
Vero cell viability assay via MTT following treatment with rfhSP-D. 5 x 10^4^ Vero cells/ well were seeded in complete MEM in 96-well culture plates and grown overnight at 37°C, 5% CO_2_. The cells were then treated with rfhSP-D (0, 10, 20, 100 μg/ml) for 24 h. 0.5 mg/ml MTT containing medium was added to the wells for 4h. The supernatants were removed, and cells were lysed using DMSO. Absorbance was measured at 590nm. Background was subtracted from all data points. The data obtained were normalised with 100% cell viability being defined as the mean of the absorbance recorded from the control sample (0 μg/ml of rfhSP-D). The data were presented as the mean of the normalised triplicates ± SEM. Significance was determined using the two-way ANOVA test and no significant reduction in cell viability was observed.

## Supplementary Raw Data

**Table S1:**
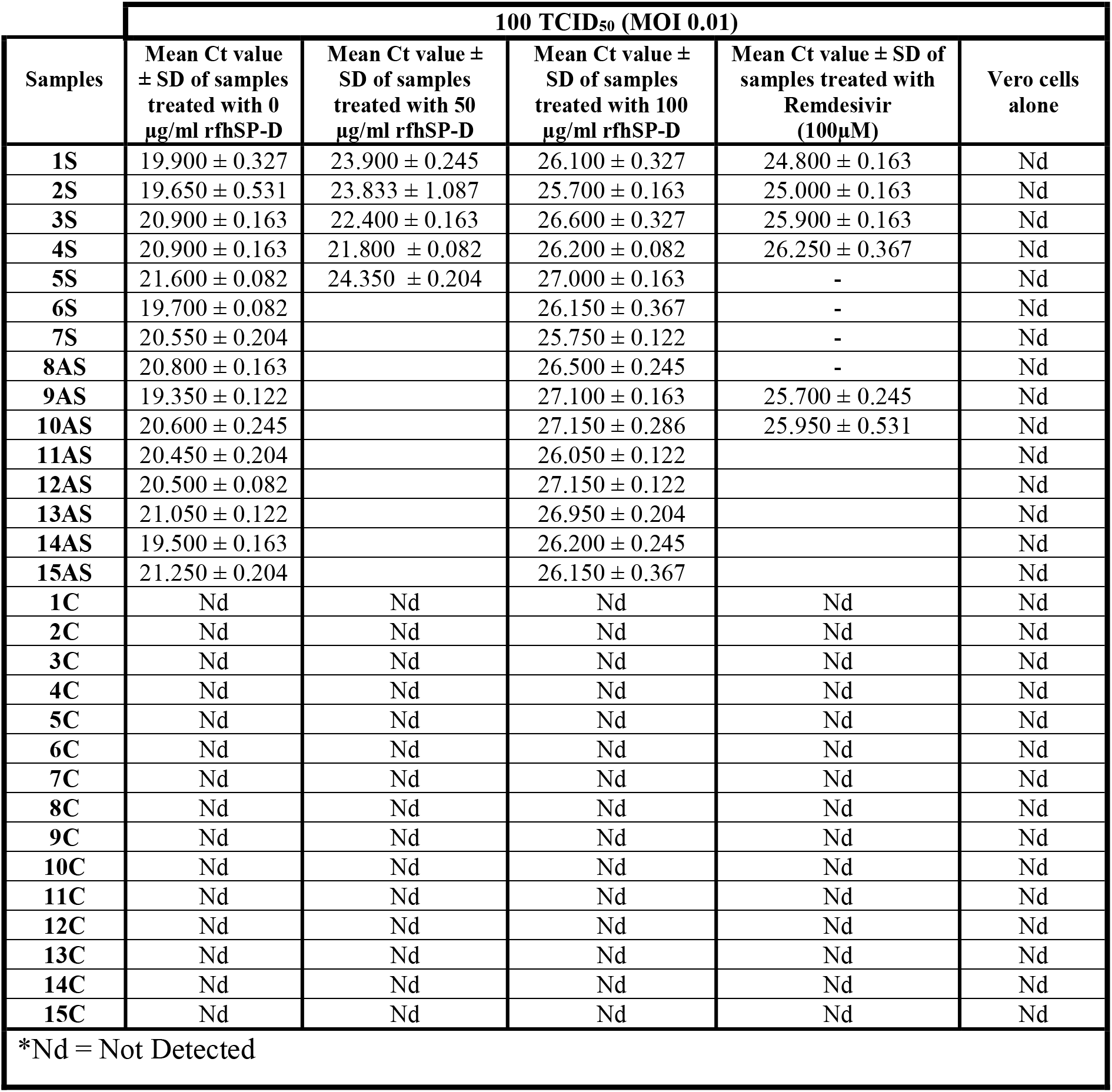
Mean Ct values of SARS-CoV-2 RdRp gene for the replication assay

**Table S2:** Mean Ct values of SARS-CoV-2 RdRp gene for the infection assay

| Sample | 500 TCID<sub>50</sub> (MOI 0.05) |  |  |  | Vero cells alone |
| --- | --- | --- | --- | --- | --- |
|  | Mean Ct value ± SD of samples treated with 0 µg/ml rfhSP-D | Mean Ct value ± SD of samples treated with 50 µg/ml rfhSP-D | Mean Ct value ± SD of samples treated with 100 µg/ml rfhSP-D | Mean Ct value ± SD of samples treated with Remdesivir (100µM) |  |
| **2S** | 21.300 ± 0.245 | 22.500 ± 0.163 | 23.400 ± 0.082 | 21.500 ± 0.163 | Nd |
| **3S** | 21.300 ± 0.327 | 22.200 ± 0.082 | 23.750 ± 0.122 | 21.600 ± 0.327 | Nd |
| **9AS** | 21.900 ± 0.163 | 23.200 ± 0.163 | 23.900 ± 0.082 | 21.900 ± 0.163 | Nd |
| **10AS** | 21.600 ± 0.163 | 22.800 ± 0.163 | 24.200 ± 0.163 | 22.200 ± 0.245 | Nd |
| **11AS** | 21.100 ± 0.163 | 23.400 ± 0.245 | 24.500 ± 0.163 | 21.850 ± 0.122 | Nd |
| **1C** | Nd | Nd | Nd | Nd | Nd |
| **2C** | Nd | Nd | Nd | Nd | Nd |
| **3C** | Nd | Nd | Nd | Nd | Nd |
| **4C** | Nd | Nd | Nd | Nd | Nd |
| *Nd = Not Detected |  |  |  |  |  |

